# A GRAPHICAL USER INTERFACE FOR WRMXPRESS 2.0 STREAMLINES HELMINTH PHENOTYPIC SCREENING

**DOI:** 10.1101/2025.03.14.643077

**Authors:** Zachary Caterer, Rachel V. Horejsi, Carly Weber, Blake Mathisen, Chase Nelson, Maggie Bagatta, Ireland Coughlin, Megan Wettstein, Ankit Kulshrestha, Hui Siang Benjamin Lee, Leonardo R. Nunn, Mostafa Zamanian, Nicolas J. Wheeler

**Affiliations:** Department of Biology, University of Wisconsin-Eau Claire, Eau Claire, WI; Department of Pathobiological Sciences, University of Wisconsin-Madison, Madison, WI

**Keywords:** anthelmintics, screening, software, imaging, phenotyping

## Abstract

Image-based phenotypic screening is a fundamental technique used to better understand the basic biology of helminths and advance discovery of new anthelmintics. Miniaturization of screening platforms and automated microscopy have led to a surge in imaging data and necessitated software to organize and analyze these data. Traditionally, these analyses are performed remotely on high-performance computers, often requiring an understanding of a command-line interface (CLI) and the ability to write scripts to control the software or job scheduler. Requiring access to specialized computing equipment and advanced computational skills raises the barrier to entry for these sorts of studies. The development of efficient, performant computer and graphical processing units for personal computers and cheaper imaging solutions has made the requirement of remote servers superfluous for many small to medium-scale screens, but most analytical software still require interaction with a CLI. To democratize the analysis of image-based phenotypic screens, we have developed a graphical user interface (GUI) for wrmXpress, a tool that integrates many popular computational pipelines for analyzing imaging data of parasitic and free-living worms. The GUI operates on any personal computer using the operating system’s native web browser, allowing users to configure and run analyses using a point-and-click approach. Containerization of the application eliminates the need to install specialized programming libraries and dependencies, further increasing the ease of use. GUI development required a substantial reorganization of the wrmXpress backend codebase, which allowed for the addition a new pipeline for high-resolution tracking of worm behavior, and we demonstrate its functionality by showing that praziquantel modulates the behavior of *Schistosoma mansoni* miracidia. These advances make cutting-edge analyses of image-based phenotyping of worms more equitable and accessible.

## 1. INTRODUCTION

The increasing demand for high-throughput and high-content imaging in biological research has required the development of efficient and user-friendly software. High-throughput imaging allows for acquisition of vast amounts of data, where high-content imaging provides detailed, multi-dimensional images of biological species (Boutros et al., 2015). Together, these technologies enable image-based phenotypic screening of parasitic roundworms and flatworms, an essential method for understanding the fundamental biology of these worms and advancing discovery of new anthelmintics (Herath et al., 2022; Zamanian and Chan, 2021).

Image-based phenotypic screening incorporates miniaturized multiwell plates, automated microscopy, and a suite of computational tools used to organize, process, and analyze the resulting data. We developed wrmXpress, an open source modular software program that allows researchers to analyze high-throughput and high-content images of parasitic flatworms and roundworms (Wheeler et al., 2022). wrmXpress, like many related tools, primarily runs remotely on high-performance computers, requiring a command-line interface (CLI) and scripting skills for software control or job scheduling. Thus, its use necessitates some informatics expertise, subsequently limiting the adoption by users who do not have computational experience or access to such collaborators. This reliance on specialized computing equipment and advanced computational skills raises the barrier to entry for many researchers.

The development of powerful new computing and graphical processing units (CPUs/GPUs) for desktop or personal computers along with affordable and/or DIY imaging options has reduced the need for remote servers for small to medium-scale screens, but most analytical software still necessitate interaction with the CLI. To democratize the analysis of image-based phenotypic screens of parasitic worms, we have redesigned wrmXpress and developed an intuitive and easy to use, point-and-click graphical user interface (GUI). This GUI preserves the full functionality of wrmXpress while bringing new capabilities to the forefront, such as a new tracking pipeline, thus enhancing the software’s scope in antiparasitic research. It also exposes method parameters to the user, allowing easy optimization to new worms and new imaging environments. Here, we present and explain the design and development of the GUI, which is containerized for deployment through a wide variety of computer platforms and accessed on common browsers, enabling access by researchers of all computational backgrounds.

## 2. METHODS

### 2.1 Design and production of the GUI

The backend of wrmXpress is primarily written in Python, which was also the language selected for developing the GUI. A variety of Python libraries exist for designing GUIs, including Dash, Tkinter, PyQt5, Kivy, and wxPython. Dash by Plotly is a framework that allows for the implementation of data visualization and user interfaces using Python, which is not often the traditional language of choice for UI and web development. Dash was selected over native and alternative graphical libraries due to its ease of use, broad user and developer base, well documented and supported code, responsive graphics, support for automated tests, and large variety of interactive visualizations while maintaining a simple yet appealing design. Core and HTML components were utilized, as well as the auxiliary Dash Bootstrap Components for styling, managing the layout, and displaying results. Dash Table was incorporated for specifying experimental metadata. Various other Python libraries we used for data handling, visualizations, image processing and displaying, YAML generation and manipulation, and testing (Bradski, 2000; Harris et al., 2020; McKinney and Others, 2010; Merkel and Others, 2014). All versioned dependencies are provided in the source code, and a frozen Anaconda environment specification file was exported with v1.0.0 of the GUI.

Creation of the GUI as a web application allows the deployment on numerous platforms (i.e., Windows, MacOS, Linux) based on user familiarity. Deployment across various systems is enabled through a pre-built Docker image that contains the frontend (GUI) and backend (wrmXpress) code (Merkel and Others, 2014; Wheeler et al., 2022). Docker facilitates consistent and reliable application usage without compromising design or efficiency, and it allows for simple synchronization of updates to the frontend or backend codebase. The app can also be run natively with Python through virtual environments, and the code is shipped with specification files for building such environments. This GUI was developed for the user with limited programming knowledge, and we recommend installing and running the app via Docker, but our approach allows the more experienced to have control over resource allocation or library versions. Documentation for installing and using the GUI can be found at https://wrmxpress-gui.readthedocs.io/latest/.

### 2.2 Parasite maintenance and harvesting

*Schistosoma mansoni* (NMRI)-infected Swiss Webster female mouse livers at 7-weeks post-infection were obtained from the Biomedical Research Institute in Rockville, Maryland, and shipped overnight in perfusion fluid (0.85% sodium chloride, 0.75% sodium citrate). Parasite eggs were harvested from the infected livers and hatched into miracidia using an adapted protocol (Dinguirard et al., 2018). Artificial pond water (APW) was prepared using the NIH NIAID Schistosomiasis Resource Center recipe (Cody et al., 2016).

Liver tissue was homogenized in 1.2% NaCl by blending with alternating speeds for 1.5 min (20 sec low/10 sec high). The homogenate was transferred to 50 mL conical tubes and washed through a series of centrifugations (500 rcf, 15 min, 4°C), until a clear supernatant was produced. After the final wash, the supernatant was decanted, and the pellet was resuspended in APW and transferred to a 1 L volumetric flask. The base of the flask was covered in aluminum foil with the top 2-4 cm left exposed to light. The flask was then filled with APW until ∼3 cm of water was above the top of the aluminum foil. Transfer pipettes were used to remove debris and bubbles at the top, and warmed APW (32-34°C) was added along the sides of the flask to create a temperature gradient. A goose-neck microscope illuminator was positioned to shine light directly onto exposed water.

After 1 hr, the top 8 mL of water, which included viable miracidia, was transferred to a 15 mL conical tube. Tubes were centrifuged for 1 min (500 rcf, 4°C) to pellet miracidia, after which excess APW was immediately removed. Miracidia were then resuspended in 1 mL of fresh APW and transferred to a 1.5 mL tube. Miracidia density was calculated by counting five 10 µL aliquots stained with 1:1 Lugol’s iodine. Miracidia were diluted with fresh APW to a final density of 1 parasite/µL.

### 2.3 Miracidia behavioral experiments

Miracidia at 1 parasite/µL were pipetted into 96-well-plates with varying treatments and their behavior was recorded using a Zeiss Stemi 508 Microscope with a DMK37BUK178 camera (The Imaging Source) and IC Capture 2.5 image acquisition software. Videos were captured using a Y800 color format, 2048×2048 field of view, focused on a region of interest of 1616×1616 pixels, and saved as uncompressed AVI files. Each video was recorded at 16 frames per second under 2X magnification. All videos of behavior were analyzed by the wrmXpress tracking and optical flow pipelines.

To evaluate the effects of praziquantel (PZQ) on *S. mansoni* miracidia, PZQ (Santa Cruz Biotechnology) was dissolved in dimethyl sulfoxide (DMSO, Santa Cruz Biotechnology) and 50X stocks were produced, yielding 500 µM, 50 µM, 5 µM, 0.5 µM, and 0.05 µM concentrations. Stocks were stored at -20°C, and the same stocks were used for all experiments. 1 µL dots of 50X PZQ were added to wells of a 96-well plate, and 49 µL of 1 hr old miracidia were added to each well. All treatments were in a final concentration of 0.1% DMSO, and this concentration of DMSO was used as a negative control. Plates were incubated for 30 min at 25°C, after which 1 min videos were recorded. Three biological replicates (miracidia hatched from separate mouse infections) were performed, with each replicate including 2-3 technical replicates (wells) per concentration for a total of N=8 for each concentration. Data was analyzed by fitting an ANOVA model for each behavioral feature.

### 2.4 Development and implementation of a miracidia tracking pipeline

Tracking was implemented into wrmXpress using the trackpy library (Allan et al., 2024). Videos are converted to grayscale and stored as a 3-dimensional Numpy array (Harris et al., 2020). The background of each video is generated by a median projection of array along the 0^th^ axis, which takes the median value of each pixel across the entire video; this approach effectively estimates the background by ignoring frames in which pixels are briefly dark when represented by a swimming worm in the foreground. This approach includes dead and unmoving worms as part of the background. Median project is slower than previously used approaches such as a rolling-mean calculation, but it is less sensitive to glare(Fogarty et al., 2022). The background is then subtracted from each frame, resulting in bright white objects (i.e., worms) on a dark background. Objects are detected with the trackpy.batch() function using empirically optimized parameters; these parameters can be adjusted in the Configuration panel of the GUI and quickly optimized using the Preview functionality. Objects are linked with trackpy.link(); objects that disappeared for >50 frames were interpreted as a new object if it later reappeared (the ‘memory’ parameter in the GUI). Linked coordinates are written out by wrmXpress as a CSV with merged metadata. All trackpy parameters can be adjusted in the Configuration panel.

Custom R scripts were used to draw tracks and extract features such as distance, velocity, tortuosity, rate of change of direction (RCD), and standard deviation of the angular velocity. All raw data and scripts can be found in a GitHub repository associated with the manuscript (https://github.com/wheelerlab-uwec/GUI_ms).

Total distance (*D*) was calculated as follows:

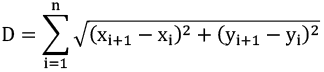

where *i* is the frame number and *n* is the total number of frames.

Velocity was calculated as total distance (*D*) divided by the total time the object was tracked. Tortuosity (τ) was calculated the total distance (*D*) divided by the chord length:

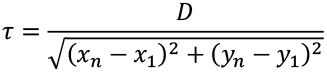

where *n* is the last frame.

RCD was calculated as the total number of directional changes in x or y divided by the total time the object was tracked.

Angular velocity (ω) was calculated as follows:

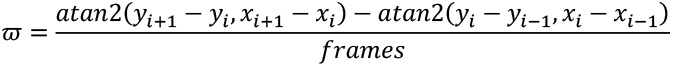

where *i* is a given frame. The standard deviation of this vector of values was calculated to generate a single value for each track.

## 3. RESULTS AND DISCUSSION

### 3.1 A graphical user interface (GUI) for wrmXpress enables phenotypic screening on personal computers

The GUI has multiple pages (Info, Configure, Metadata, Preview, and Run), each with a specific and intuitive purpose. The Info page consists of information regarding the design and implementation of the GUI and the backend code and some instructions regarding usage of the application. More explicit usage instructions and reference material can be found at the associated documentation site (wrmxpress-gui.readthedocs.io/latest/). Additional information such as proper file formatting and information for who to contact for support is also included.

As originally designed, wrmXpress options are configured with a user-provided YAML file, and the GUI Configure page enables the simple creation of this strictly formatted file based on the user selections. The Configure page contains numerous features allowing for the user to understand their selections prior to analysis, including tooltips that display brief descriptions of given buttons or inputs, previews of pipeline outputs, and automated dynamic updates of configuration options based on user input (Fig. 1A-C). The page also enables the selection of specific wells to analyze, allowing a user to save time by only running the pipeline on key wells of interest (Fig. 1C). Primary purposes of the GUI are to mitigate configuration errors, allow for optimization of parameters, and enable simple job initiation rather than requiring command line programming to deploy jobs to a remote server. The Configure page simplifies pipeline and parameter selection and dynamically warns users of missing parameters or selections that are likely to lead to errors during analysis. Redesign of the wrmXpress backend allowed for the exposure of nearly all analytical or algorithmic parameters to the GUI which, used in tandem with the Preview functionality (see below), allows for the efficient optimization of parameters.

**Figure 1.**
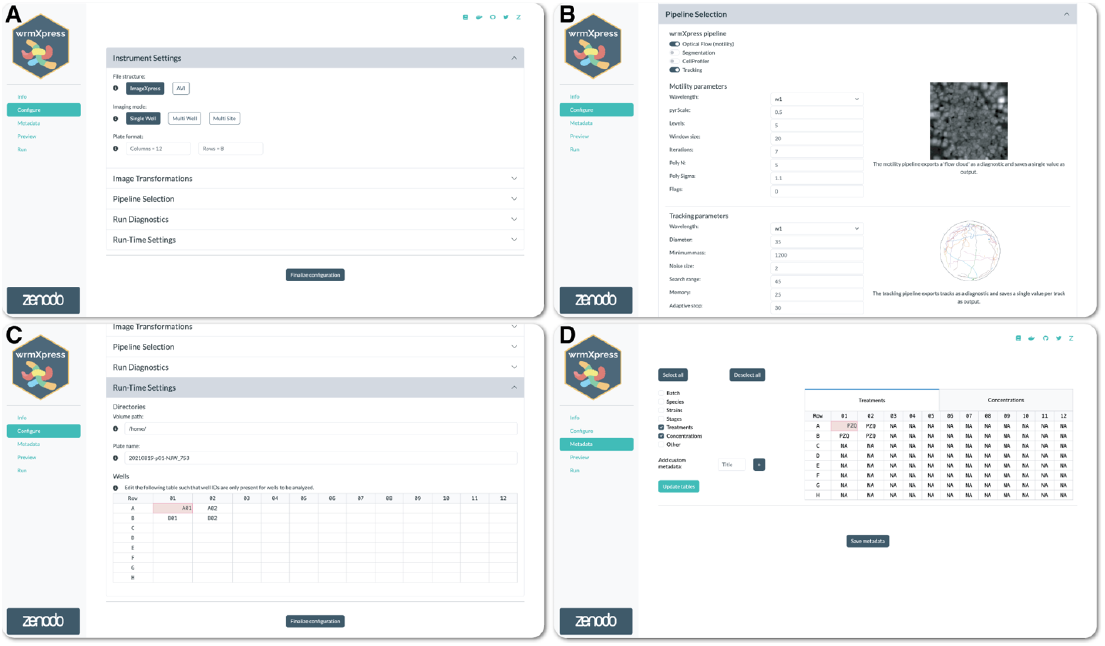
Configuration of a wrmXpress analysis. (A) Selections for instrument settings. (B) Pipeline selection along with example output image and description. Multiple pipelines can be run on the same input dataset. (C) Path configuration and well selection. In this case, the Volume path directs to a mounted volume in a Docker container. Four wells are selected for analysis. (D) Well-based metadata input, which is flexible and allows for custom metadata entries. NAs are automatically generated for easy post hoc filtration.

Configuration parameters are stored internally and accessible to other pages. For instance, the Metadata page allows the user to input custom metadata, which are then saved as CSV files and joined as new columns to the output tabular data generated by the pipeline (Fig. 1D). The structure of the Metadata input table is determined by selections on the Configure page (i.e., the number of rows and columns in the plate to be analyzed).

After pipeline and parameter selection, analysis of the input images or videos is performed through the Preview and Run pages. The Preview page offers a new way to anticipate results of the user’s selections by conducting the analysis on a single well rather than all selected wells (Fig. 2). This information provides the user with informative expected results, allowing for refinement of parameters prior to conducting the full analysis. Once the user has finalized the configuration and verified these selections in the Preview page, they can conduct the full analysis using the Run page. Each pipeline is processed in a similar well-based iteration, and a progress bar is dynamically updated during run-time (Fig. 3A). Updates to the wrmXpress backend also now allow for multi-site imaging, and the user can analyze each site independently or stitch sites together prior to analysis. Independent analysis allows for input derived from 384-well plates, which often tile wells in a 2x2 image rather than 384 separate images.

**Figure 2.**
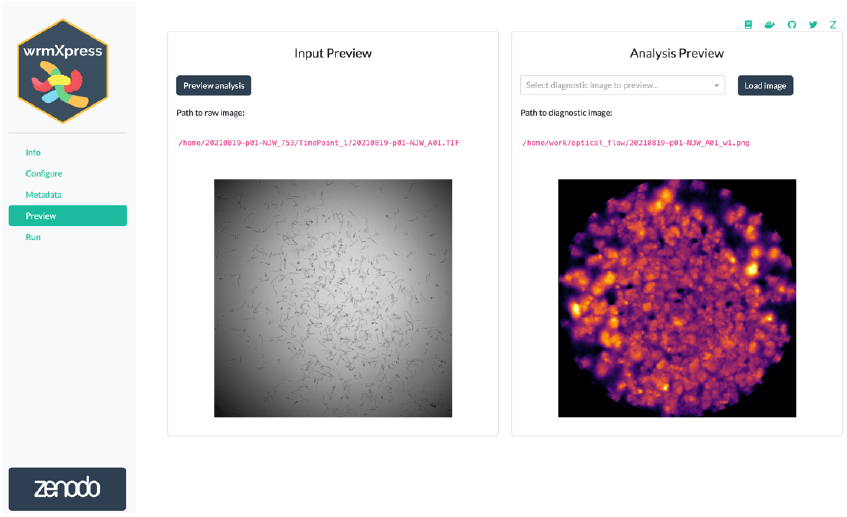
Previewing a wrmXpress run. The GUI will run the configured analysis on the first of the selected wells and show a diagnostic image for the selected pipeline. The path to the preview diagnostic is provided for further inspection. If multiple pipelines are selected, preview diagnostics are generated for each. A “flow cloud” diagnostic of worm motility is shown.

**Figure 3.**
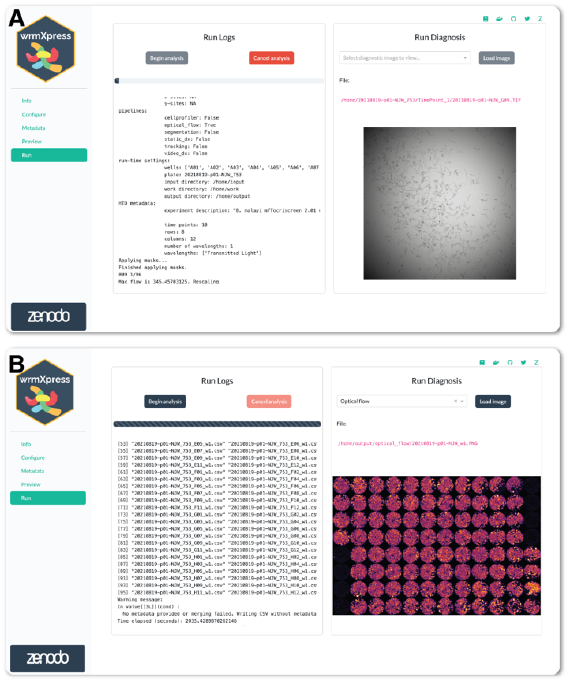
Running wrmXpress. (A) Log messages are written to the screen during a run and a progress bar is updated based on the number of selected wells. A thumbnail of the well being analyzed is shown and dynamically updated. Log files are also written to the disk for later inspection or troubleshooting. (B) Well-based diagnostic images are stitched together once analysis is complete to provide an overview of the entire plate. In the example plate analyzed with the motility pipeline, black wells were positive control wells filled with heat-killed worms.

While running wrmXpress, log messages are printed to the Run page. Logs are also written to a file, facilitating bug and issue reports that can be delivered to the application maintainers. At run completion, wells from the entire analysis are stitched together to provide a set of final diagnostic images for the plate (Fig. 3B). Pipelines tested within the GUI on 8 wells of previously generated data completed in 3-70 seconds per well (Table 1), which is a reasonable amount of time to expect for image-based analyses on a personal computer.

**Table 1.** Pipeline run times when run in a Docker container (https://hub.docker.com/t/wheelern/wrmxpress_gui) with 32 GB allocated RAM and 14 CPUs using an Apple MacBook Pro M3 Max running MacOS Sonoma 14.5. Eight wells from test data were analyzed in three separate runs. *Docker was not used, but the GUI was run natively using a Conda environment to allow for full utilization of the machine’s RAM/CPUs.

| *Pipeline* | *Wells* | *Mean Run-Time (s)* | *Standard Deviation (s)* |
| --- | --- | --- | --- |
| *Worm size (Cellpose (Stringer et al., 2021))* | 8 | 539.07 | 2.63 |
| *Worm size* | 8 | 240.44 | 0.82 |
| *Fecundity* | 8 | 24.77 | 0.02 |
| *Feeding* | 8 | 220.95 | 0.42 |
| *Motility* | 8 | 66.87 | 0.05 |
| *Viability* | 8 | 81.07 | 0.07 |
| *Tracking\** | 8 | 783.48 | 0.61 |

### 3.2 The wrmXpress tracking pipeline creates full-featured datasets for individual worms and reveals specific effects of praziquantel on the behavior of schistosome miracidia

In addition to a new graphical frontend for wrmXpress, further updates to the backend were made. A tracking pipeline was implemented, which tracks the movement of individual worms by generating frame-by-frame coordinate data. The tracking pipeline requires an amount of RAM that scales linearly with the number and resolution of video frames. Long videos with a high resolution may require RAM that is more than what is available on an individual computer or allocatable to a Docker container. This is demonstrated by the longer runtime for the tracking pipeline when compared with other pipelines (Table 1).

We demonstrated the utility of this pipeline by using it to analyze the effects of praziquantel (PZQ) on *Schistosoma mansoni* miracidia, the aquatic snail-infective stage of trematodes. Previously, it was shown that 3 µM PZQ can totally inhibit the motility of freshly hatched miracidia after 1 hr or 1 mM after 5 min (Liang et al., 2001; Ryan et al., 2023). In these experiments, effectiveness of PZQ was essentially binary – either miracidia were swimming, or they were not. We expected that tracking the behavior of individual worms would reveal more sensitive effects of PZQ.

In general, 1 hr old miracidia will swim straight in a helical motion with periodic bouts of increased turning. When treated with PZQ, behavioral differences are clearly observable in tracks generated by the new pipeline (Fig. 4A). Increasing concentration of PZQ within individual wells resulted in reduced movement of miracidia (Fig. 4A-B).

**Figure 4.**
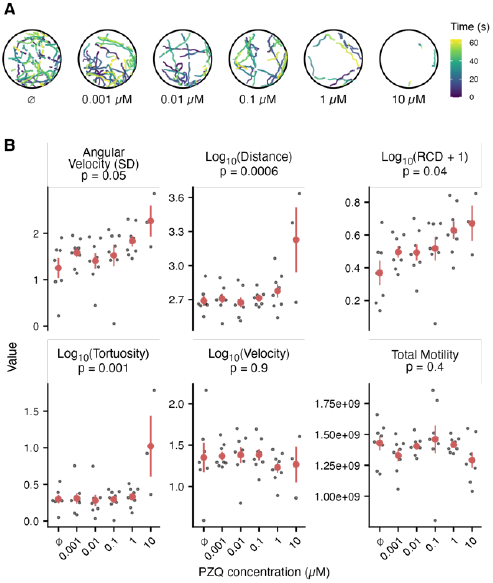
Praziquantel (PZQ) affects the swimming behaviors of *Schistosoma mansoni* miracidia. (A) Sample tracks of miracidia after 30 min. incubation with different concentrations of PZQ. Videos were recorded for 1 min. The color of track represents the time of recording in seconds. (B) Behavioral features of miracidia after treatment with PZQ. Each black point represents the mean value of each well; red points and ranges represent the mean and standard error of the mean. p-values were calculated by fitting an ANOVA model.

In many other parasitic worm species and stages, optical flow as a measure of total motility is routinely implemented (Marcellino et al., 2012; Storey et al., 2014), and wrmXpress incorporates this approach in its motility pipeline (Wheeler et al., 2022). However, for most of these worms, translational movement in liquid media is limited, and thrashing is the primary neuromuscular output. In contrast, miracidia are native to liquid culture and show directional movement. We found that optical flow was not an informative approach for measuring drug effects for miracidia, perhaps because this difference in motility. This is likely the reason for binary measures of drug effects on miracidia in the past (Liang et al., 2001; Ryan et al., 2023). In contrast, our new tracking pipeline measures different behavioral characteristics that are more sensitive and informative than total motility as measured with optical flow (Fig. 4B). An ANOVA analysis revealed significant differences within the behavioral characteristics newly produced by the pipeline (p < 0.05). The lack of significant differences of total motility due to PZQ treatment gives us reason to believe that optical flow does not work well for tracking miracidia although it has been successful when used with other worms. Tracking miracidia through the new pipeline allows for the extraction of behavioral features in response to different treatments. Overall, these results indicate the functionality of this new tracking pipeline and its potential to advance the field of miracidia ethology and anthelminthic discovery.

## 4. CONCLUSIONS

The wrmXpress GUI significantly improves the accessibility of cutting-edge image-based phenotypic screening of worms to researchers without extensive computational backgrounds. This intuitive, easy to use, and interactive GUI operates on any personal computer through its native web browser, removing the need for specialized software and preexisting computational knowledge and skills. The GUI integrates seamlessly with the current wrmXpress software, preserving all previous functionality while also adding the tracking pipeline, which allows for the sensitive analysis of worm behavior, especially those that show directed translation in liquid culture. The GUI is fully documented, further reducing the barrier to entry.

The modular nature of the GUI and the redesign of the backend facilitates expansion and updating, ensuring it can maintain relevance and meet future needs of researchers and technology advancements. Such future advancements may consist of development of additional pipelines, adoption of machine learning and artificial intelligence, or extension beyond the realm of parasitological research. Indeed, many pipelines could be adapted for use with other aquatic invertebrates. Through this GUI we have democratized access to advanced phenotypic screening techniques, promoting a more inclusive environment, encouraging collaboration and innovation for scientists around the world.

Installation instructions for the wrmXpress GUI can be found at the documentation website (https://wrmxpress-gui.readthedocs.io/). The prebuilt Docker image can be found at Docker Hub (https://hub.docker.com/r/wheelern/wrmxpress_gui). Example data for running each pipeline can be found on Zenodo (https://doi.org/10.5281/zenodo.12760651). The wrmXpress GUI is distributed under the CC BY-NC-SA 4.0 license.

## 5. ACKNOWLEDGEMENTS

*S. mansoni*-infected mouse tissue was provided by the Schistosomiasis Resource Center of the Biomedical Research Institute (Rockville, MD) through NIH-NIAID Contract HHSN272201700014I. Funding was provided by Student Blugold Commitment Differential Tuition funds through the University of Wisconsin-Eau Claire and the UWEC Office of Academic Affairs. Some computational resources used for this study were provided by the Blugold Center for High-Performance Computing under NSF grant CNS-1920220. MZ was supported by NIH NIAID R01 AI151171. NJW was supported by NIH NIAID R15 AI183095. The authors would like to thank, Katie Ryan and Natalia Betancourt for training Cellpose models, members of the Zamanian and Wheeler labs for assistance with editing of the final manuscript, and Tom and Joe Berg for beta testing and assistance with CSS styling.

